# Bioenergetic stress potentiates antimicrobial resistance and persistence

**DOI:** 10.1101/2024.07.12.603336

**Authors:** B Li, S Srivastava, M Shaikh, G Mereddy, MR Garcia, A Shah, N Ofori-Anyinam, T Chu, N Cheney, JH Yang

## Abstract

Antimicrobial resistance (AMR) is a global health crisis and there is an urgent need to better understand AMR mechanisms. Antibiotic treatment alters several aspects of bacterial physiology, including increased ATP utilization, carbon metabolism, and reactive oxygen species (ROS) formation. However, how the “bioenergetic stress” induced by increased ATP utilization affects treatment outcomes is unknown. Here we utilized a synthetic biology approach to study the direct effects of bioenergetic stress on antibiotic efficacy. We engineered a genetic system that constitutively hydrolyzes ATP or NADH in *Escherichia* coli. We found that bioenergetic stress potentiates AMR evolution via enhanced ROS production, mutagenic break repair, and transcription-coupled repair. We also find that bioenergetic stress potentiates antimicrobial persistence via potentiated stringent response activation. We propose a unifying model that antibiotic-induced antimicrobial resistance and persistence is caused by antibiotic-induced. This has important implications for preventing or curbing the spread of AMR infections.

## INTRODUCTION

Antimicrobial resistance (AMR) is a global health crisis and poses a looming threat to modern medicine. Antimicrobial resistant infections are estimated to have been associated with 4.95 million global deaths and have directly caused 1.27 million global deaths in 2019^1^. There is therefore an urgent need to better understand AMR mechanisms.

AMR is defined by heritable protection from antimicrobial agents. Resistance is most frequently caused by genetic acquisition of mutations that inhibit drug-target interactions or alter drug transport^2^. Non-heritable forms of phenotypic resistance also exist, including antimicrobial persistence^3^. Persistence is caused by stochastic formation of an isogenic sub-population of “persister cells” that are insensitive to antimicrobial stress^4–6^. Persistence can enable downstream resistance evolution^7–9^.

Mechanisms underlying antibiotic-induced resistance and persistence are poorly understood^10,11^. Growing evidence demonstrate antibiotic treatment exerts pleiotropic stresses that potentiate mutation rates thereby accelerating AMR evolution (stress-induced mutagenesis)^8,9,12–17^. These stresses amplify several aspects of bacterial physiology including ATP utilization^18–21^, cellular respiration^22,23^, and production of reactive species (ROS)^21,22,24^. Imbalanced ATP utilization and production creates “bioenergetic stress”, which impairs growth and enhances glycolysis, oxidative phosphorylation, and ROS formation^25–29^. ROS stimulate stress-induced mutagenesis by multiple DNA repair mechanisms^8,12^ and molecules which inhibit these processes decelerate AMR evolution^14,15^.

Antimicrobial persistence is associated with down-regulated metabolic activity and decreased intracellular ATP^4,30–35^. Genetic studies implicate stress-induced activation of the (p)ppGpp-mediated stringent response as an important component to persister cell formation^6,34,36^. However, specific mechanisms linking antimicrobial stress to stringent response activation remain unclear.

While antibiotic-induced ROS formation is now well studied^10,11,24,37^, much less is known on how antibiotic-induced bioenergetic stress regulates antibiotic treatment responses. Here we utilized a synthetic biology approach to directly study how bioenergetic stress impacts antimicrobial resistance and persistence in *Escherichia coli*. We found that bioenergetic stress alone can accelerate fluoroquinolone resistance evolution via a stress-induced mutagenesis mechanism involving ROS, mutagenic break repair, and transcription-coupled repair. We found that bioenergetic stress also potentiates persister cell formation via stringent response and that the stringent response contributes to resistance evolution. Together, our results provide a unifying model for how antibiotic stress can enhance antimicrobial resistance and persistence by altering bacterial physiology.

## RESULTS

### Bioenergetic stress potentiates antibiotic resistance evolution and persistence

Our previous work implicate antibiotic-induced ATP utilization as an inducer of hyper-respiratory activity^18^. To determine if antibiotic treatment induces bioenergetic stress, we metabolically profiled antibiotic-treated *E. coli* MG1655 cells by LC-MS/MS^38,39^ (Supplementary Table 1). One hour treatment with 16 ng/mL of the fluoroquinolone ciprofloxacin (2x MIC: minimum inhibitory concentration) significantly decreased abundance of the bioenergetic metabolites ATP and NADH, resulting in decreased ATP/ADP and decreased NADH/NAD^+^ ratios (Fig. 1a).

**Figure 1.**
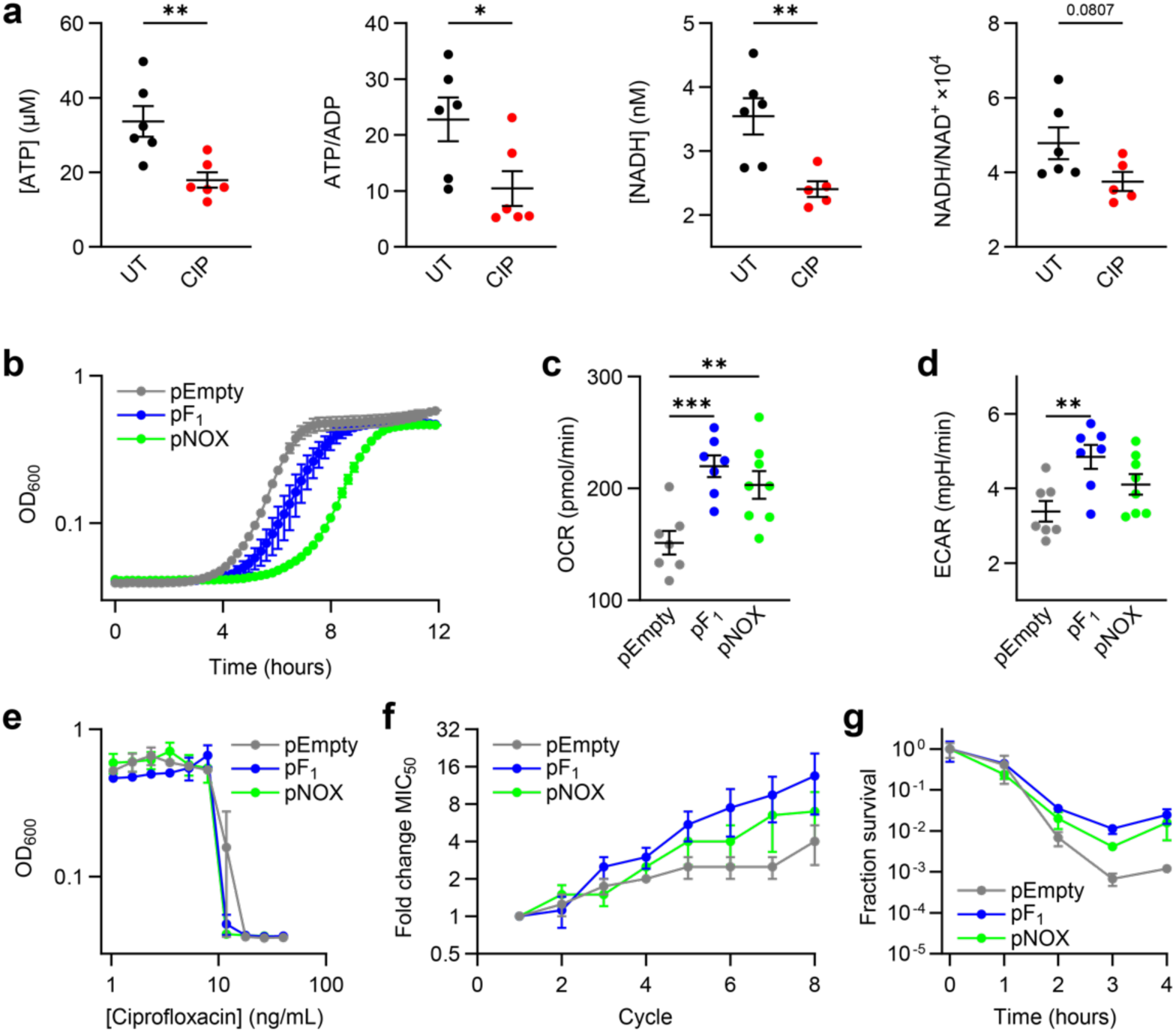
Bioenergetic stress potentiates antimicrobial resistance evolution and persistence. **a**, Energetic metabolite profiles for untreated (UT) and ciprofloxacin treated (CIP) *E. coli* MG1655 cells as determined by LC-MS/MS. Exponential phase cells were grown in MOPS minimal media and treated ± 16 ng/mL ciprofloxacin for 1 hour (n = 6). **b**, Growth curves for pEmpty, pF_1_, and pNOX cells grown in MOPS rich media. **c**, Exponential phase oxygen consumption rates (OCR) as a reporter of respiratory activity. **d**, Exponential phase extracellular acidification rates (ECAR) as a reporter of glycolytic activity. **e**, Growth measurements in the presence of ciprofloxacin after 1:20,000 dilution from stationary phase. **f**, Changes in ciprofloxacin susceptibility over 8 cycles of serial-passage laboratory evolution. Data reported as change in the minimum concentration for 50% growth inhibition (MIC_50_) relative to Cycle 1. **g**, Ciprofloxacin lethality following treatment with 18 ng/mL ciprofloxacin. Data reported as change in colony forming units (CFUs) relative to time 0. Data depicted as mean ± 95% CI. All experiments performed in MOPS rich media unless otherwise indicated. n = 4 for all experiments unless otherwise indicated. All data depicted as mean ± SEM unless otherwise indicated. \**p* ≤ 0.05, \*\**p* ≤ 0.01, \*\*\**p* ≤ 0.001. Non-significant comparisons not shown.

To directly investigate the effects of bioenergetic stress on antibiotic efficacy without the potential confounding effects of changes in bacterial physiology induced by antibiotic treatment, we took a synthetic biology approach. We engineered a genetic system comprised of constitutive over-expression of *E. coli*’s soluble ATP synthase F_1_ complex (*atpAGD*, pF_1_) or constitutive heterologous expression of the NADH oxidase from *Streptococcus pneumoniae* (*nox*, pNOX) on low-copy plasmids^25–29^. pF_1_ or pNOX induce bioenergetic stress by continuously hydrolyzing ATP or NADH, respectively. This hydrolysis constitutively increases ATP or NADH utilization and demand, thus futilely cycling ATP or NADH production. Consistent with previous reports, constitutive expression of these enzymes impaired fitness (Fig. 1b), increased respiration (Fig. 1c and Extended Data Fig. 1a; increased OCR: oxygen consumption rate), and increased glycolysis (Fig. 1d and Extended Data Fig. 1b; increased ECAR: extracellular acidification rate), when compared with empty vector controls (pEmpty). To further characterize this system, we measured single-cell changes in membrane polarization by fluorescence microscopy using the membrane potential-sensitive probe DiOC_2_(3). We found significantly decreased membrane potential in pNOX but not pF_1_ expressing cells (Extended Data Fig. 1c-e). This is consistent with expectation as increased glycolysis is expected to increase respiratory Complex I activity and Complex I can function as a H^+^/Na^+^ antiporter in *E. coli*^40–42^. Together, these data validated our genetic system.

We next evaluated the effects of bioenergetic effects of antibiotic susceptibility. pF_1_ or pNOX expression did not increase the MIC for ciprofloxacin, indicating bioenergetic stress does not directly confer genetic resistance (Fig. 1e). However, serial passage laboratory evolution experiments revealed unexpected acceleration of ciprofloxacin resistance evolution (Fig. 1f). Moreover, time-kill experiments revealed unexpected increases in the fraction of persister cells surviving ciprofloxacin treatment (Fig. 1g). Collectively, these data indicate bioenergetic stress alone can enhance both genotypic and phenotypic AMR.

### Bioenergetic stress induces oxidative DNA damage

To better understand how bioenergetic stress alters bacterial physiology, we sequenced RNA from exponential phase pEmpty, pF_1_, and pNOX cells. Importantly, sequencing analysis revealed increased expression of *atpA*, *atpG*, and *atpD* in only pF_1_ and *nox* in only pNOX cells (Extended Data Fig. 2a-c).

Statistical analyses revealed only 72 differentially expressed genes between pF_1_ and pEmpty cells (≥ 2-fold change, FDR-corrected p ≤ 0.05) and only 314 differentially expressed genes between pNOX and pEmpty cells (Supplementary Table 2). Bioinformatic analyses of the 72 shared genes with altered expression between pF_1_ and pNOX cells revealed only unexpected upregulation of genes involved in chemotaxis and flagellar assembly and downregulation of genes involved in hydrogen sulfide metabolism (Supplementary Table 3 and Extended Data Fig. 2d). Genetic deletion of CheY or FlhD, the transcription factors regulating chemotaxis or flagellar assembly gene expression, did not reverse the increased persistence caused by bioenergetic stress (Extended Data Fig. 2e). Bioinformatic analyses further revealed no changes in the basal induction of genes involved in DNA repair, stringent response, or oxidative stress response processes, which are implicated as mechanisms of antimicrobial resistance and persistence. The decreased basal expression of H_2_S biosynthesis genes was surprising as H_2_S protects against bactericidal antibiotics^43,44^, indicating bioenergetically induced persistence was also likely not due to hydrogen sulfide production.

Because bioenergetic stress alters bacterial metabolism^25–29^, we performed genome-scale metabolic modeling analyses^45^ to predict how bioenergetic stress would globally remodel *E. coli* metabolism. We applied our RNA sequencing data as modeling constraints^46,47^ to the comprehensive iML1515 model of *E. coli* metabolism^48^ and performed flux variability analysis^49^. Consistent with our experimental measurements, model simulations predicted increased ATP synthase activity, NADH production, oxygen consumption, and hexokinase (first step in glycolysis) activity in both pF_1_ and pNOX cells over pEmpty cells (Fig. 2a and 2b). These results validated our metabolic modeling simulations.

**Figure 2.**
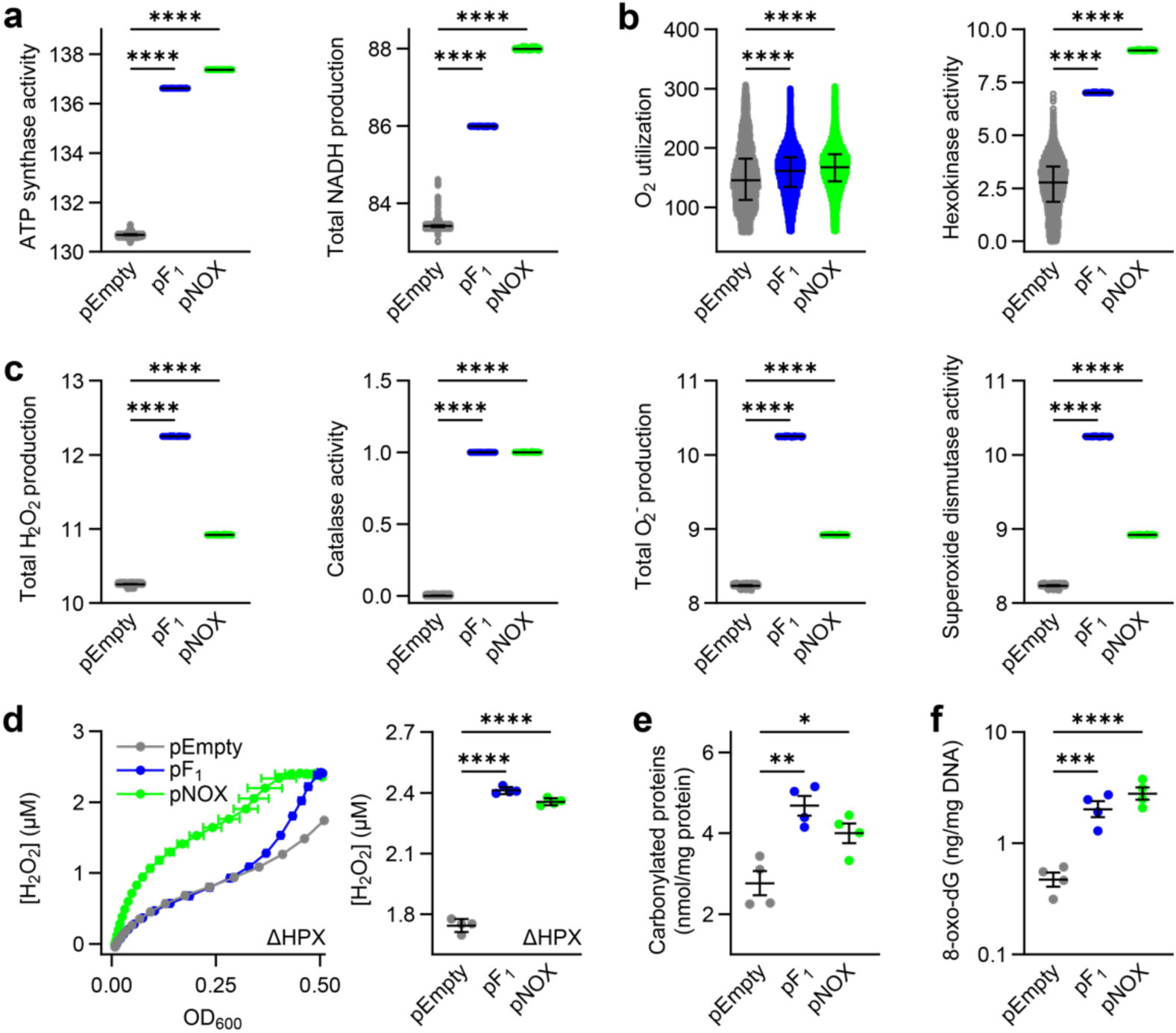
Bioenergetic stress induces oxidative cellular damage. **a**, ATP and NADH production rates predicted by flux variability analyses (FVA) using the iML1515 genome-scale model of *E. coli* metabolism^48^ (n ≥ 10,000 flux samples for all model simulations). **b**, Predicted respiratory (oxygen utilization) and glycolytic (hexokinase) activities. **c**, Predicted intracellular H_2_O_2_ production, catalase activity, superoxide production, and superoxide dismutase activity rates. Error bars depict median flux distributions ± interquartile ranges. **d**, Integrated H_2_O_2_ production by catalase and peroxidase-deficient Δ*ahpCF* Δ*katE* Δ*katG* (ΔHPX) cells expressing pEmpty, pF_1_, or pNOX (*left*). Integrated H_2_O_2_ at mid-exponential phase (OD_600_ ≈ 0.5; *right*). **e**, Carbonylated proteins in exponential phase pEmpty, pF_1_, and pNOX cells. **f**, Deoxyguanosine oxidation in exponential phase pEmpty, pF_1_, and pNOX cells. All experiments performed in MOPS rich media. n = 4 for all experiments. All data depicted as mean ± SEM unless otherwise indicated. \**p* ≤ 0.05, \*\**p* ≤ 0.01, \*\*\**p* ≤ 0.001, \*\*\*\**p* ≤ 0.0001. Non-significant comparisons not shown.

Although the expression of oxidative stress response regulators *oxyR*, *oxyS*, *soxR*, or *soxS* did not increase (Supplementary Table 3 and Extended Data Fig. 3), model simulations predicted significant increases in basal H_2_O_2_ production, catalase activity, superoxide production, and superoxide dismutase activity (Fig. 2c). These results suggested pF_1_ and pNOX expression increased basal ROS production. To test this hypothesis, we measured H_2_O_2_ production in each strain using a highly specific and quantitative Amplex UltraRed horseradish peroxidase assay in a catalase and peroxidase-deficient Δ*ahpCF* Δ*katG* Δ*katE* (ΔHPX) background^50^ (Fig. 2d). Consistent with previous observations that ATP futile cycling enhances H_2_O_2_ production^28^, we observed increased basal H_2_O_2_ production in both ΔHPX pF_1_ and ΔHPX pNOX cells. Because we did not observe significant induction of OxyR or SoxRS response regulators, we hypothesized the increased ROS production would result in oxidative cellular damage. To test this hypothesis, we performed enzyme-linked immunosorbent assays (ELISAs) for carbonylated proteins and 8-oxo-deoxyguanosine and found increased basal oxidative damage to proteins (Fig. 2e) and DNA (Fig. 2f). These results are interesting because they reveal that bacteria can undergo oxidative stress that is sufficient for cellular damage but is undetected by oxidative stress responses^51^. Collectively, these results indicate bioenergetic stress induces basal oxidative DNA damage.

### Bioenergetic stress accelerates resistance evolution via enhanced ROS production

Deoxyguanosine oxidation is highly mutagenic^52^. We therefore hypothesized that bioenergetic stress accelerates resistance evolution by potentiating mutagenesis. To test this hypothesis, we performed Luria-Delbruck fluctuation assays^53,54^ to quantify basal mutation rates. Surprisingly, basal mutation rates did not differ between pF_1_, pNOX, and pEmpty cells, suggesting that bioenergetic stress accelerated resistance evolution by enhancing stress-induced mutagenesis (Extended Fig. 4a).

We hypothesized that the physiological changes induced by bioenergetic stress (increased respiration and ROS production) were responsible for the accelerated ciprofloxacin resistance evolution. To test this hypothesis, we utilized both genetic and biochemical approaches. We previously demonstrated that the respiratory mutants Δ*atpA* or Δ*cyoA ΔcydB ΔappB* (ΔETC) increased or decreased respiratory activity, respectively (Fig. 3a and Extended Fig. 4b). We found *atpA* deletion to also enhance H_2_O_2_ production (Fig. 3b and Extended Fig. 4c). These mutants are deficient for functional ATP synthase or cytochrome oxidase complexes, respectively. We performed ELISAs for carbonylated proteins and 8-oxo-deoxyguanosine and found that Δ*atpA* cells also induced oxidative cellular damage to levels like catalase and peroxidase-deficient ΔHPX cells (Fig. 3c and 3d). These results demonstrate increased respiration is sufficient for increasing ROS production.

**Figure 3.**
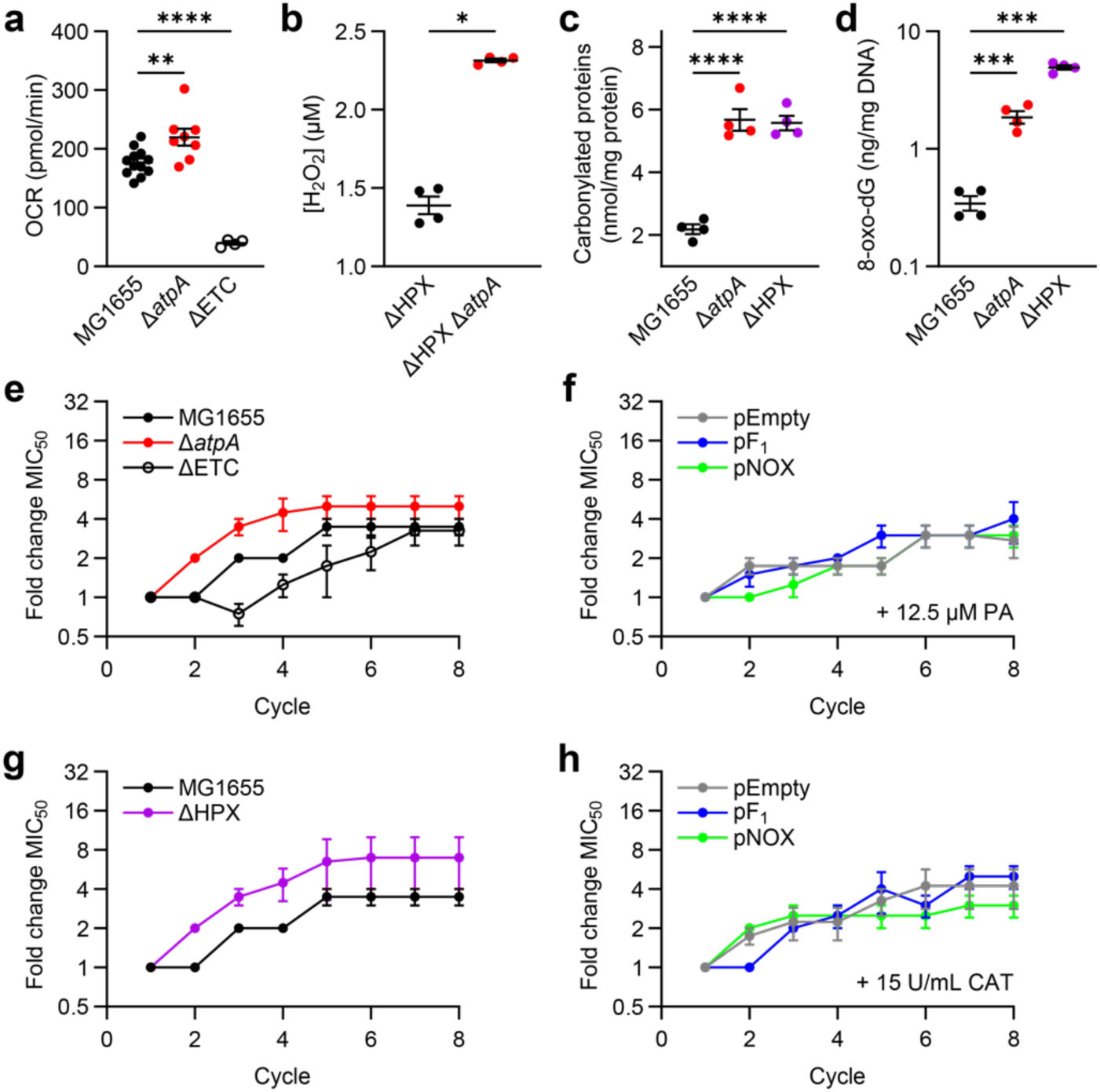
Bioenergetic stress accelerates resistance evolution by enhancing ROS production. **a**. Exponential phase oxygen consumption rates (OCR) for wild-type MG1655, Δ*atpA*, and Δ*cyoA* Δ*cydB* Δ*appB* (ΔETC) *E. coli* cells (n ≥ 4). **b**. Integrated H_2_O_2_ for ΔHPX and ΔHPX Δ*atpA* cells at mid-exponential phase (OD_600_ ≈ 0.5). **c**, Carbonylated proteins in exponential phase MG1655, Δ*atpA*, and ΔHPX cells. **d**, Deoxyguanosine oxidation in exponential phase MG1655, Δ*atpA*, and ΔHPX cells. **e**, Ciprofloxacin resistance evolution for MG1655, Δ*atpA*, and ΔETC cells. **f**, Ciprofloxacin resistance evolution for pEmpty, pF_1_, and pNOX cells in the presence of 12.5 µM respiratory inhibitor piceatannol (PA). **g**, Ciprofloxacin resistance evolution for MG1655 and ΔHPX cells. **h**, Ciprofloxacin resistance evolution for pEmpty, pF_1_, and pNOX cells in the presence of 15 U/mL catalase (CAT). Resistance evolution data reported as change in the minimum concentration for 50% growth inhibition (MIC_50_) relative to Cycle 1 for all resistance evolution experiments. All experiments performed in MOPS rich media. n = 4 for all experiments unless otherwise indicated. All data depicted as mean ± SEM unless otherwise indicated. \**p* ≤ 0.05, \*\**p* ≤ 0.01, \*\*\**p* ≤ 0.001, \*\*\*\**p* ≤ 0.0001.

To determine if respiration and ROS regulate AMR evolution, we performed laboratory evolution experiments with Δ*atpA* and ΔETC cells. Ciprofloxacin resistance evolution was accelerated in high-respiring Δ*atpA* cells, but not in low-respiring ΔETC cells (Fig. 3e). To determine if differences in respiratory activity were responsible for the accelerated resistance evolution induced by pF_1_ or pNOX expression, we repeated the ciprofloxacin resistance evolution experiments in the presence of 12.5 μM piceatannol supplementation and observed no differences between pF_1_, pNOX, and pEmpty cells (Fig. 3f). Piceatannol is an ATP synthase inhibitor^55^ that decreases cellular respiration in *E. coli* without inhibiting growth (Extended Data Fig. 4d-f). These results indicate increased respiration contributes to bioenergetic-stress accelerated resistance evolution.

To determine if differences in oxidative stress were responsible for the accelerated resistance evolution induced by pF_1_ or pNOX expression, we performed similar experiments in ΔHPX cells (Fig. 3g) or in the presence and absence of 15 U/mL catalase (Fig. 3h). We found resistance evolution to be accelerated in ΔHPX cells and found no differences in resistance evolution between pF_1_, pNOX, and pEmpty cells. Together, these results demonstrate that bioenergetic stress potentiates antimicrobial resistance evolution via increased respiration and increased ROS.

### Bioenergetic stress induces Type 1 persistence

We next sought to understand how bioenergetic stress potentiates antimicrobial persistence. To determine if persistence, like accelerated resistance, was caused by increased respiration and ROS, we performed time kill experiments in the presence of supplemented piceatannol or catalase. Similar to low-respiring ΔETC cells (Fig. 4a), respiratory inhibition by piceatannol inhibited ciprofloxacin lethality (Fig. 4b)^23^. ROS detoxification by catalase did not prevent the increased persistence observed in pF_1_ or pNOX cells (Fig. 4c). These demonstrated that the observed increases in resistance and persistence caused by bioenergetic stress involved different mechanisms.

**Figure 4.**
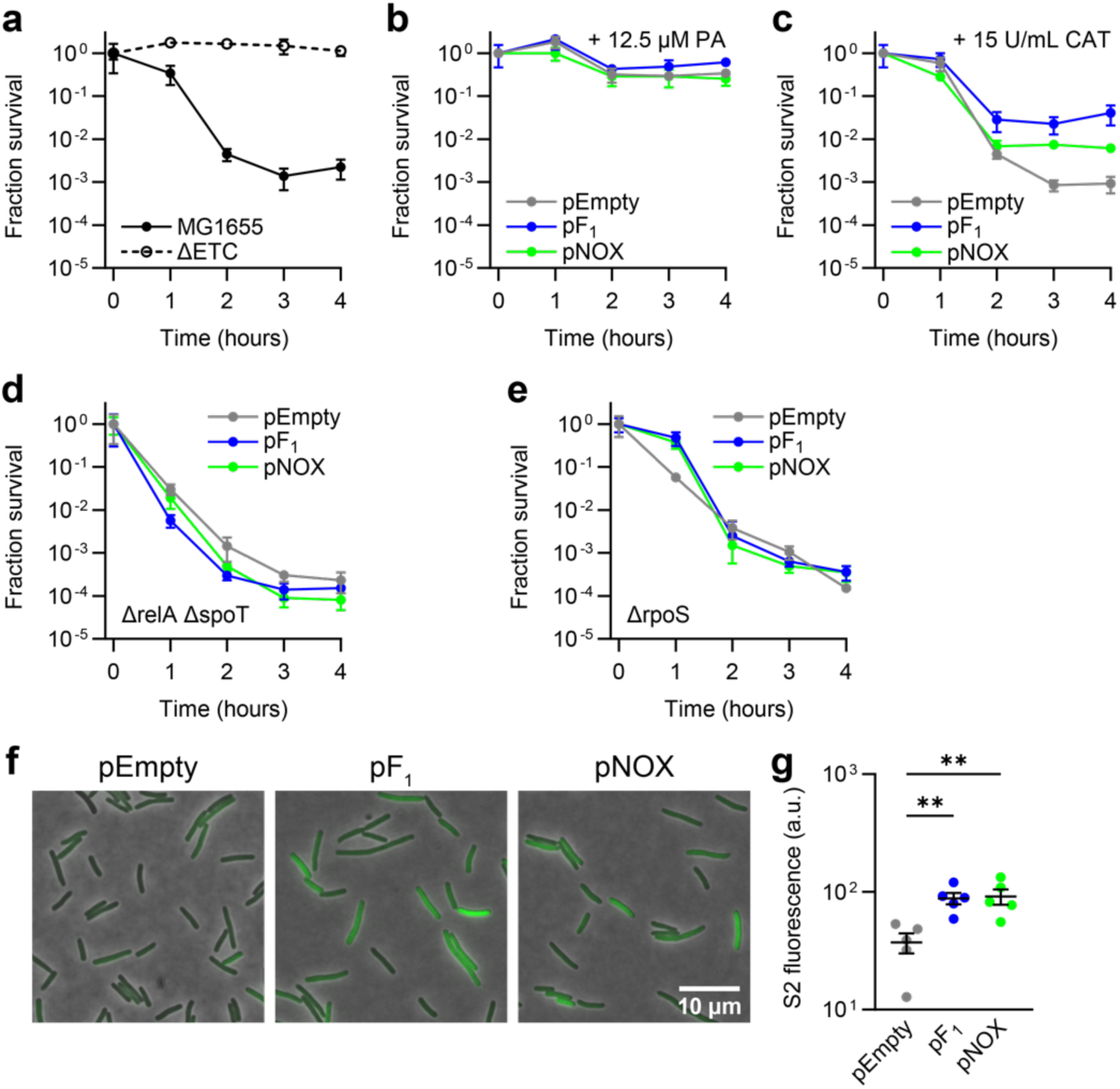
Bioenergetic stress potentiates persistence via enhanced stringent response activation. **a**, Ciprofloxacin lethality following treatment with 16 ng/mL ciprofloxacin for wild-type MG1655 and ΔETC cells. **b**, Ciprofloxacin lethality following treatment with 18 ng/mL ciprofloxacin for pEmpty, pF_1_, and pNOX cells in the presence of 12.5 µM piceatannol (PA). **c**, Ciprofloxacin lethality following treatment with 18 ng/mL ciprofloxacin for pEmpty, pF_1_, and pNOX cells in the presence of 15 U/mL catalase (CAT). **d**, Ciprofloxacin lethality following treatment with 18 ng/mL ciprofloxacin for Δ*relA ΔspoT* cells expressing pEmpty, pF_1_, or pNOX. **e**, Ciprofloxacin lethality following treatment with 18 ng/mL ciprofloxacin for Δ*rpoS* cells expressing pEmpty, pF_1_, or pNOX. Data reported as change in colony forming units (CFUs) relative to time 0 for all time-kill experiments. Data depicted as mean ± 95% CI. **f**, Representative fluorescence microscopy images for exponential phase *E. coli* BL21(DE3) cells expressing the S2 ppGpp fluorescent biosensor^58^ and pEmpty, pF_1_, or pNOX after 1 hour treatment with 18 ng/mL ciprofloxacin. **g**, S2 fluorescence quantification (n = 5). Data points depict medians of mean cell intensities. Error bars depict grand mean ± SEM from n ≥ 135 cells per replicate. All experiments performed in MOPS rich media. n = 4 for all experiments unless otherwise indicated. \*\**p* ≤ 0.01.

Persister cell formation frequently involves activation of the (p)ppGpp-mediated stringent response which ubiquitously responds to diverse stressors by inducing growth inhibition and metabolic dormancy^56^. Stringent response activation involves synthesis of the alarmone (p)ppGpp by the pyrophosphokinase RelA and the (p)ppGpp hydrolase SpoT, which binds and activates the transcription factor DksA, inducing several pleiotropic effects including activation of the general stress response sigma factor RpoS. To determine if bioenergetic stress potentiates persistence by enhancing stringent response activation, we performed time-kill experiments in Δ*relA ΔspoT* (Fig. 4d) or *ΔrpoS* (Fig. 4e) cells expressing pF_1_, pNOX, or pEmpty. These experiments revealed no differences in persistence between pEmpty and pF_1_ or pNOX cells, implicating stringent response activation as the mechanism for bioenergetic-stress potentiated persistence.

Persister cell formation can be either environmentally triggered (Type 1) or stochastically formed during exponential growth (Type 2)^57^. To determine if bioenergetic stress induced Type 1 or Type 2 persistence, we performed fluorescence microscopy experiments using the S2 genetically encoded (p)ppGpp fluorescent biosensor^58^. S2 fluorescence was higher in pF_1_ and pNOX cells than pEmpty cells when treated with 18 ng/mL ciprofloxacin for 1 hour (Fig. 4f and 4g). Importantly, S2 fluorescence was not different between pF_1_, pNOX, or pEmpty during exponential growth (Extended Data Fig. 5a and 5b). Collectively, these results demonstrate that bioenergetic stress induces Type 1 persistence.

### Bioenergetic stress-induced persistence inhibits respiration-induced lethality

Our findings that persistence is increased in high-respiring pF_1_ and pNOX cells seemingly contradict our previous findings that increased respiration enhances antibiotic killing^23^. Indeed, high-respiring Δ*atpA* cells exhibited growth defects, increased respiration, and increased glycolytic activity like bioenergetically stressed cells (Fig. 3a and Extended Data Fig. 4b, 6a, and 6b), but displayed faster lethality and lower persistence than wild-type cells (Fig. 5a). Because Δ*atpA* cells lack functional ATP synthase and share metabolic physiology with pF_1_ and pNOX cells, we hypothesized that Δ*atpA* cells would similarly be bioenergetically stressed. To test this hypothesis, we metabolically profiled Δ*atpA* cells by LC-MS/MS and surprisingly did not observe decreases in ATP or NADH ratios that would be indicative of bioenergetic stress (Fig. 5b, Extended Data Fig. 6c and 6d, and Supplementary Table 5).

**Fig. 5:**
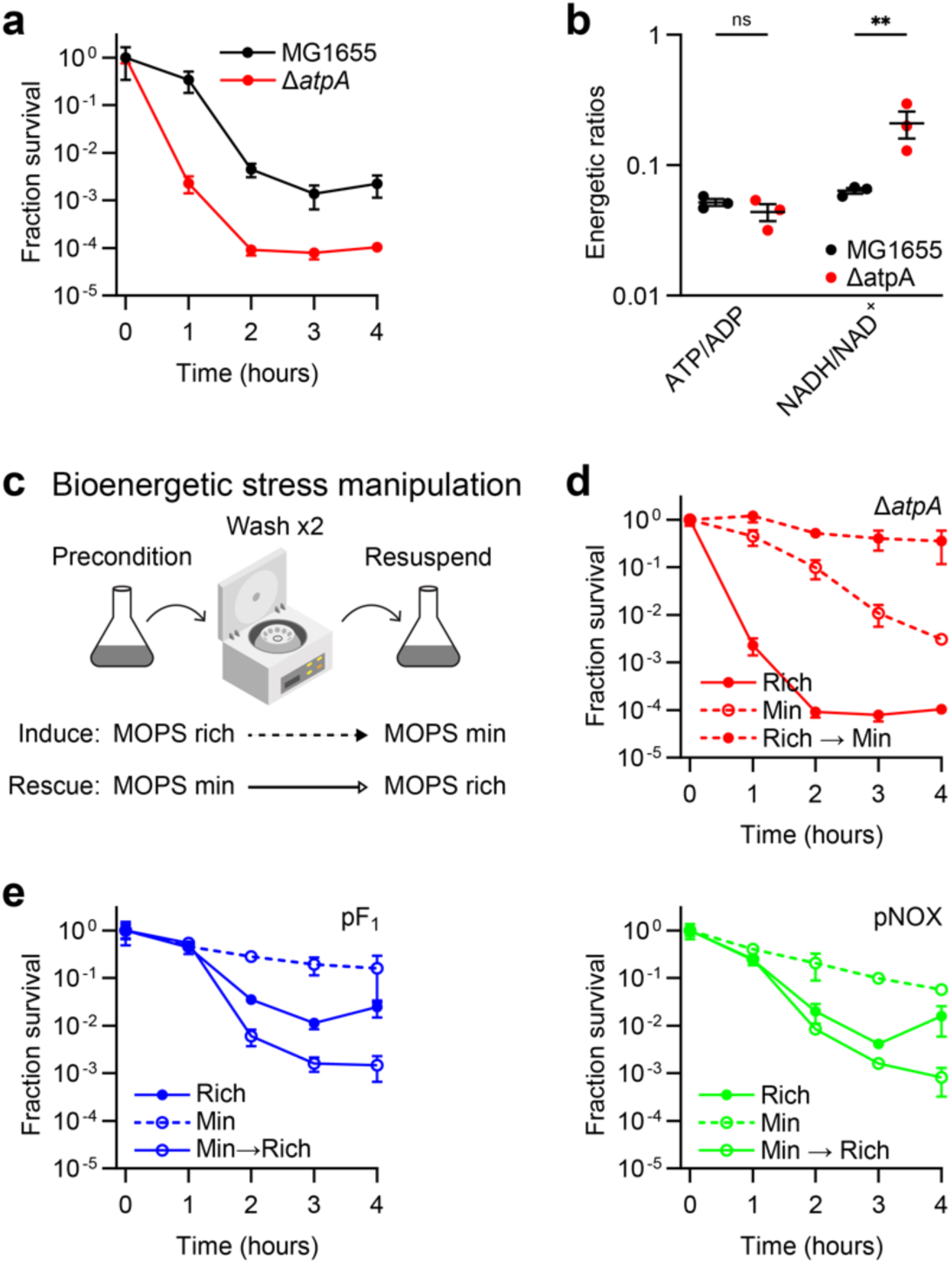
Bioenergetic stress is required for stress-induced persistence. **a**, Ciprofloxacin lethality following treatment with 16 ng/mL ciprofloxacin for wild-type MG1655 and Δ*atpA* cells in MOPS rich media. **b**, Energetic ratios for exponential phase MG1655 and Δ*atpA* cells grown in MOPS minimal media (n = 3). **c**, Schematic for bioenergetic stress manipulation by media switching. Cells were bioenergetically preconditioned by growth in MOPS rich or minimal media. Exponential phase cells were washed twice in PBS and then resuspended in rich or minimal media containing ciprofloxacin. Cultured were sampled hourly and plated for CFU enumeration after media switching. Bioenergetic stress is induced by resuspending cells preconditioned in rich media into minimal media. Bioenergetic stress is rescued by resuspending cells preconditioned in minimal media into rich media. **d**, Ciprofloxacin lethality for Δ*atpA* cells grown and treated with 16 ng/mL ciprofloxacin in rich media, minimal media, or following a switch from rich media to minimal media. Bioenergetic stress induces persistence in Δ*atpA* cells. **e**, Ciprofloxacin lethality for pF_1_ or pNOX cells grown and treated with 18 ng/mL ciprofloxacin in rich media, minimal media, or following a switch from minimal media to rich media. Bioenergetic rescue decreases persistence in pF_1_ and pNOX cells. n = 4 for all experiments unless otherwise indicated. Data reported as change in colony forming units (CFUs) relative to time 0 for all time-kill experiments. Data depicted as mean ± 95% CI for all time-kill experiments. ns > 0.05, \*\**p* ≤ 0.01.

These results support the hypothesis that bioenergetic stress specifically induces persister cells that are protected from antibiotic lethality. To further test this hypothesis, we performed media-changing experiments to manipulate bioenergetic stress in high respiring cells (Fig. 5c). In non-stressed cells, ATP production and protein expression is matched to ATP utilization to establish energetic equilibrium. The rationale for this experiment is as follows. When ATP utilization is rapidly altered, ATP production and ATP utilization are transiently unbalanced until protein expression and cellular metabolism is remodeled to restore homeostasis. Because ATP utilization is higher under growth in minimal media than rich media^59^, a rapid switch from rich media to minimal media causes ATP utilization to transiently exceed ATP production (thereby inducing bioenergetic stress), while a rapid switch from minimal media to rich media induces the opposite effect (thereby rescuing bioenergetic stress).

We performed time kill experiments in Δ*atpA* cells where exponential phase cells grown in rich media were rapidly switched to minimal media containing ciprofloxacin. We found that bioenergetically stressed Δ*atpA* cells were significantly protected from ciprofloxacin (Fig. 5d). Importantly, this protection exceeded the decreased lethality observed in Δ*atpA* cells grown in minimal media indicating the change in lethality was not due to only the extracellular minimal media environment. Consistent with these results, we found that recuing bioenergetic stress in pF_1_ or pNOX cells by switching exponential phase cells grown in minimal media to rich media containing ciprofloxacin increased ciprofloxacin lethality when compared with cells grown and treated in minimal media (Fig. 5e). Importantly, this increase in lethality also exceeded the level of killing observed in cells grown and treated in rich media.

Together, these results demonstrate that the persistence induced by bioenergetic stress can overcome the lethality imposed by high respiration and high ROS.

### Bioenergetic stress accelerates resistance evolution via “gambler cell” formation and transcription-coupled repair

We next sought to understand the DNA repair mechanisms underlying bioenergetic stress-potentiated resistance. Two mechanisms were recently proposed for ROS-enhanced ciprofloxacin resistance evolution. Rosenberg et al. propose that ROS potentiates ciprofloxacin resistance evolution by enabling a sub-population of hyper-mutagenic “gambler” cells^8^. This mechanism involves stringent response activation^9^ and low-fidelity DNA polymerase IV (*dinB*). Merrikh et al. propose that ROS-induced DNA damage induces stalled RNA polymerases, which recruit the DNA translocase Mfd to initiate nucleotide excision repair of oxidative DNA lesions via UvrABC^12,13^.

To test the involvement of gambler cells, we performed ciprofloxacin resistance evolution experiments in *ΔdinB* or *ΔrelA ΔspoT* cells expressing pF_1_, pNOX, or pEmpty. We did not find any differences in resistance evolution between *ΔdinB* pF_1_, pNOX, and pEmpty cells (Fig. 6a), nor differences in *ΔrelA ΔspoT* pF_1_ and pEmpty cells (Fig. 6b). These results supported the hypothesis that bioenergetic stress potentiates gambler cell formation.

**Figure 6.**
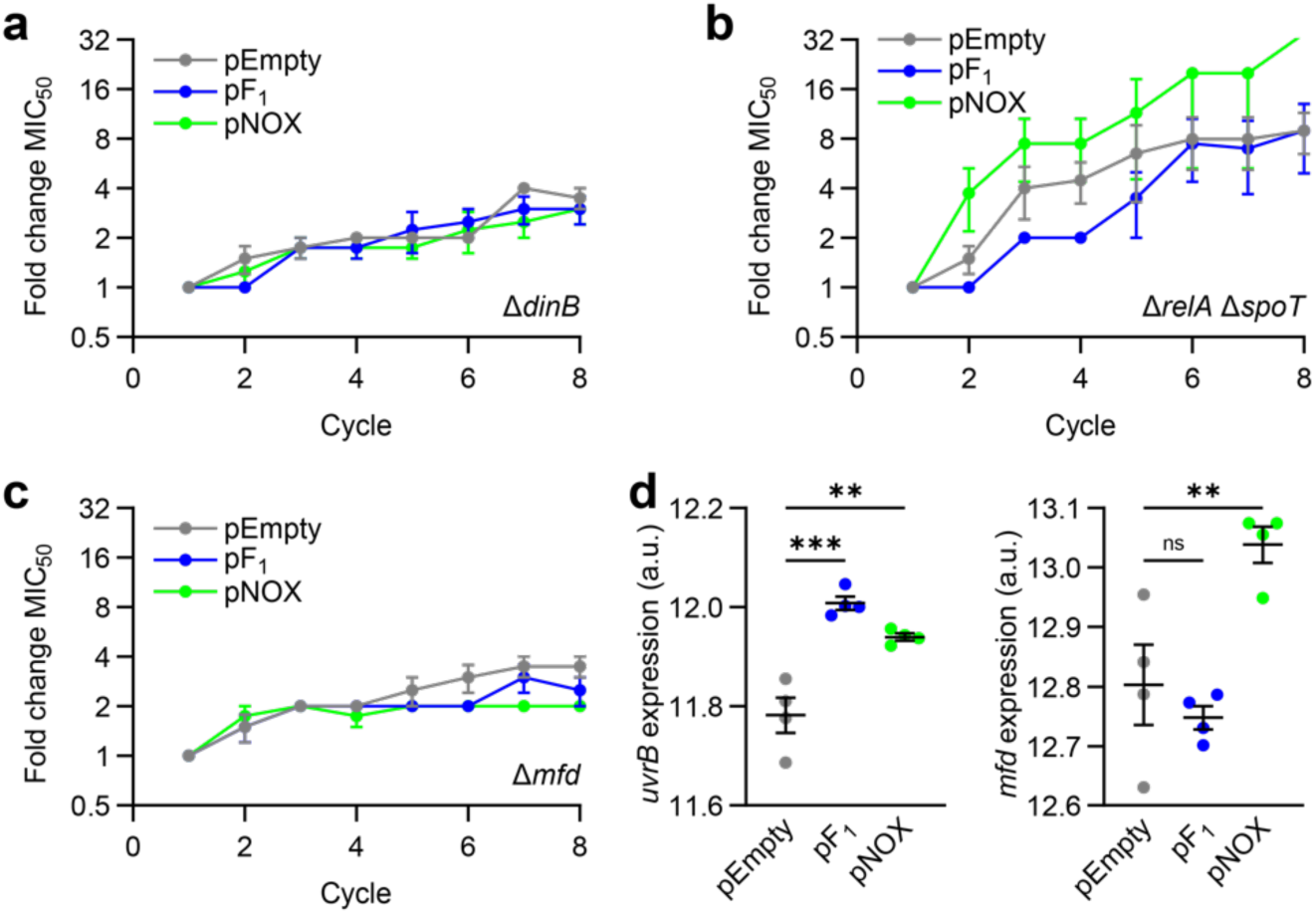
Bioenergetic stress accelerates antimicrobial resistance evolution by both mutagenic break repair and transcription-coupled repair. **a**, Ciprofloxacin resistance evolution for *ΔdinB* cells expressing pEmpty, pF_1_, or pNOX. **b**, Ciprofloxacin resistance evolution for *ΔrelA ΔspoT* cells expressing pEmpty, pF_1_, or pNOX. **c**, Ciprofloxacin resistance evolution for *ΔrelA ΔspoT* cells expressing pEmpty, pF_1_, or pNOX. Data reported as change in the minimum concentration for 50% growth inhibition (MIC_50_) relative to Cycle 1 for all resistance evolution experiments. **d**, *uvrA* and *mfd* expression for exponential phase pEmpty, pF_1_, and pNOX cells. Data depicted as quantile normalized log_2_-transformed transcripts per million counts as measured by RNA-sequencing. All experiments performed in MOPS rich media. n = 4 for all experiments. ns > 0.05, \*\**p* ≤ 0.01, \*\*\**p* ≤ 0.001.

To test the involvement of transcription-coupled repair, we performed ciprofloxacin resistance evolution experiments in *Δmfd* cells expressing pF_1_, pNOX, or pEmpty. We did not find any differences in resistance evolution between these cells (Fig. 6c). These results also supported the hypothesis that bioenergetic stress enhances Mfd-dependent mutagenesis. In further support of this hypothesis, we observed modestly increased *uvrB* and *mfd* expression in pF_1_ and pNOX cells over pEmpty cells (Fig. 6d).

Thus, our data suggest bioenergetic stress accelerates resistance evolution by both gambler cell formation and transcription-coupled repair through DNA repair mechanisms beyond the scope of this study.

## DISCUSSION

Our findings here reveal several unexpected and important insights into how bioenergetic stress impacts antimicrobial resistance, persistence, and bacterial physiology (Fig. 7). Despite commonly held notions on how metabolic dormancy protects against antimicrobial stress^4,30–35^, our data demonstrate metabolic quiescence is not required for persistence. Instead, our data suggest metabolic persistence is caused by imbalances between ATP utilization and production, which can occur when ATP utilization increases (e.g., during antimicrobial stress^18–20)^ or when ATP production decreases (e.g., in stationary phase or under nutrient limitation^60^). This is likely a general phenomenon not restricted to antibiotic treatment or metabolic dormancy, as many conditions associated with persistence also induce bioenergetic stress, including acute hypoxia^61^, oxidative stress^62^, and acid stress^63^.

**Figure 7.**
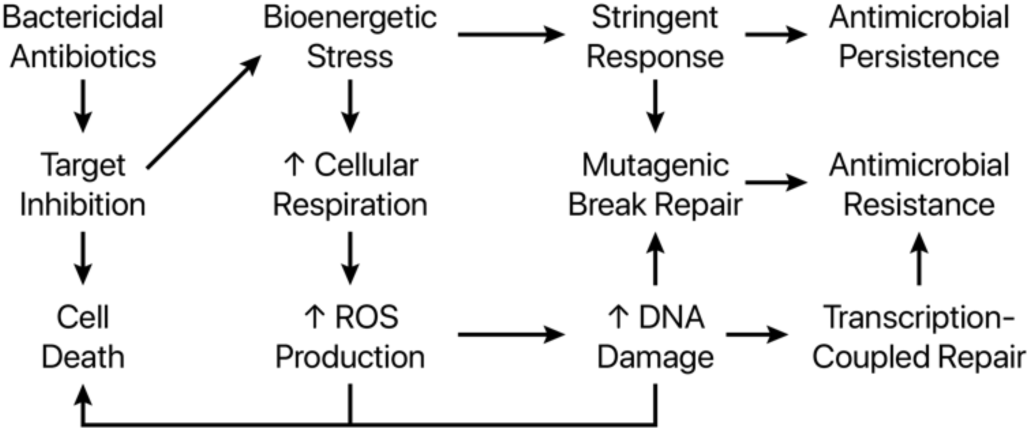
Antibiotic-induced bioenergetic stress potentiates antimicrobial resistance and persistence by altering bacterial physiology. Antibiotic treatment induces bioenergetic stress by increasing ATP utilization^52,53^. Bioenergetic stress increases cellular respiration and ROS production. Increased ROS accelerates resistance evolution by mutagenic break repair and transcription-coupled repair. Bioenergetic stress potentiates antimicrobial persistence by enhancing antibiotic-induced stringent response activation. Stringent response activation accelerates resistance evolution by mutagenic break repair.

Our data also advances understanding into the relationship between cellular respiration and antimicrobial efficacy. Our data show that although respiration and ROS potentiate antibiotic lethality^10,11,23,24,37,64^, bioenergetic stress-induced activation of the stringent response can overcome these potentiating effects, even in high-respiring or oxidatively stressed cells. These results may explain some of the misunderstandings on the role of ROS in antibiotic lethality^65–67^.

How bioenergetic stress mechanistically induces the stringent response remains unknown. The (p)ppGpp synthetases RelA and SpoT are biochemically activated by only uncharged tRNAs or acyl carrier proteins, respectively^56^. These are likely pleiotropically regulated by cellular bioenergetics, but this has not yet been directly shown.

Moreover, the mechanisms by which antibiotic treatment and ROS induces and potentiates resistance evolution is an area of active investigation^8,9,12,13,68–71^. Our data support the hypothesis that both mutagenic break repair and transcription-coupled repair^8,9^ participate in bioenergetic stress-induced resistance evolution^12,13^. Our findings that bioenergetic stress-potentiated resistance evolution is curbed in Δ*relA ΔspoT* cells is consistent with findings by others that the stringent response is involved in antimicrobial resistance evolution^71–75^. Our data support the hypothesis that persister cells can facilitate resistance evolution^7^ by providing a reservoir of hyper-mutagenic “gambler cells”^9^. However, other mutagenic DNA repair mechanisms are likely involved. Such mechanisms include base-excision repair^76^ and mismatch repair^77^ pathways that are also involved in ROS-induced mutagenesis. Understanding these mechanisms will enable the exciting development of anti-evolution drug adjuvants that can prevent or curb the expansion of drug-resistant infections^13–17,78^.

Finally, our findings have several translational implications for research domains beyond antimicrobial resistance. Heterologous expression of genes and proteins imposes bioenergetic stresses that create significant barriers for metabolic engineering and synthetic biology^79–81^. Understanding the sources of bioenergetic stress, physiological responses to bioenergetic stress, and physiological consequences of bioenergetic stress will enable development of microbiological interventions that will enhance the robustness of biochemical production in bioreactors and the stability of gene circuits in synthetic biology applications. Indeed, because bioenergetic metabolites such as ATP and NADH are universally essential cofactors across the tree of life^82^, much is to be explored in understanding the mechanisms and physiological consequences of bioenergetic stress.

## METHODS

### Bacterial strains, media, growth conditions, and chemical reagents

All experiments were performed in *Escherichia coli.* Strains used in this study were wild-type MG1655, MG1655 Δ*cheY*, MG1655 Δ*flhD*, MG1655 Δ*ahpCF ΔkatE ΔkatG* (ΔHPX), MG1655 Δ*atpA*, ΔHPX Δ*atpA*, MG1655 Δ*cyoA ΔcydB ΔappB* (ΔETC), MG1655 Δ*dinB*, MG1655 Δ*mfd*, MG1655 Δ*relA ΔspoT*::kan^R^, MG1655 Δ*rpoS*, and pET-28c-S2 expressing BL21(DE3) strains expressing pEmpty, pF_1_, or pNOX. Details on strains used in this study are provided in Supplemental Table 6.

Unless otherwise specified, cells were cultured in MOPS EZ rich defined media (Teknova; Hollister, CA). For cloning, cells were cultured in Luria-Bertani (LB) broth. For metabolomics experiments, cells were cultured in MOPS EZ minimal media (Teknova). Cells were either grown at 37°C in baffled flasks or 14 mL test tubes with 300 rpm shaking, on in round bottom 96-well plates with 900 rpm shaking, or in an Infinite M Plex plate reader (Tecan; Mannedorf, Switzerland) with 900 rpm shaking.

All antibiotics and chemical reagents were purchased from Sigma-Aldrich (St. Louis, MO) unless otherwise specified. Uniformly labeled ^13^C glucose was purchased from Cambridge Isotope Laboratories (Tewksbury, MA). Strains with antibiotic selection markers were grown in media containing 30 μg/mL chloramphenicol, 100 μg/mL ampicillin, or 50 μg/mL kanamycin. All experiments were performed with n ≥ 3 from independent overnight cultures starting from single colonies on LB agar plates (BD; Franklin Lakes, NJ).

### Gene knockout strain construction

MG1655 Δ*atpA*, Δ*cheY*, Δ*flhD*, Δ*dinB*, Δ*mfd*, and Δ*rpoS* strains were constructed by P1 phage transduction using the Keio collection^83^, as described previously^18^. Briefly, overnight cultures of *E. coli* BW25113 kan^R^ cells grown in LB media were inoculated 1:100 into fresh media supplemented with 0.2% glucose and 5 mM CaCl_2_ and allowed to grow for 1 hour before P1 phage was added. After 2 hours of incubation, phage lysates were passed through a 0.22 μm filter to remove remaining bacteria. An overnight culture of MG1655 was pelleted by centrifugation at 4,200 rcf for 10 minutes and resuspended in a 10 mM MgCl_2_ and 5 mM CaCl_2_ salt solution, then incubated with the phage lysate at 37°C for 30 minutes. LB containing 5 mM sodium citrate was added to each tube and incubated at 37°C for an additional 60 minutes in a 300 rpm shaking incubator. Cells were pelleted by centrifugation at 6,000 rcf for 2 minutes, resuspended in fresh media containing 5 mM sodium citrate, then plated on kanamycin-selective LB agar plates containing 5 mM sodium citrate. After 24 hours of incubation, colonies were selected from each plate and their kanamycin-resistance cassettes were cured by transforming the pCP20 plasmid^84^ via electroporation, inducing recombination by overnight growth at 42°C, and then screening resulting colonies for genomic recombination and plasmid loss on kanamycin- and ampicillin-selective LB agar plates. ΔHPX was constructed by curing the kanamycin-resistance cassette from MG1655 Δ*ahpCF ΔkatE*::kan^R^ Δ*katG*^83^ by pCP20 as described above. ΔHPX Δ*atpA* was constructed by P1 phage transduction of Δ*atpA*::kan^R^ into ΔHPX and curing the kanamycin-resistance cassette as described above.

### Plasmid generation

The plasmid backbone shared by pEmpty, pF_1_, and pNOX consists of a p15 origin of replication, chloramphenicol resistance, and a constitutive synthetic promoter derived from pAB191:lacZ^85^. pEmpty was generated by PCR amplification and restriction cloning of the plasmid backbone without the lacZ gene insert. pF_1_ was generated by PCR amplification of *atpAGD* from *E. coli* MG1655, PCR amplification of the synthetic promoter from pCP41::atpAGD plasmid^27^, and by restriction cloning of these fragments into pEmpty. pNOX was generated by PCR amplification of the fragment containing the synthetic promoter and *Streptococcus pneumoniae* NADH oxidase from the pAC06::nox plasmid^47^ and by restriction cloning into pEmpty. All constructs were fully sequenced for validation. Plasmid maps are available on request.

### Metabolomic characterization

Intracellular metabolites from ciprofloxacin-treated or untreated *E. coli* cells were extracted and quantified on an AB SCIEX Qtrap 5500 mass spectrometer (AB SCIEX; Framingham, MA) as described previously^38,39^. Briefly, overnight cultures of MG1655 cells grown in MOPS minimal media were inoculated 1:500 into fresh media and grown until they reached OD_600_ ≥ 0.1. Cells were back-diluted to OD_600_ = 0.1 in 25 mL MOPS minimal media, dispensed into 250 mL baffled flasks, and treated with 16 ng/mL ciprofloxacin or solvent control (H_2_O). Samples were collected 1 hour after supplementation, and aliquots with biomass equivalents to 10 mL of cell culture at OD_600_ = 0.1 were subjected to metabolite extraction using a 40:40:20 mixture of acetonitrile, methanol, and LC-MS grade water. Uniformly labeled ^13^C-standards were generated by growing *E. coli* in uniformly labeled glucose M9 minimal media in baffled flasks, as described previously. Calibration standards were split across several mixes, aliquoted, and lyophilized to dryness. All samples and calibrators were equally spiked with the same internal standards. Samples were quantified using isotope-dependent mass spectrometry. Calibration curves were run before and after all biological and analytical replicates. Consistency of quantification between calibration curves was checked by running a Quality Control sample composed of all biological replicates. Intracellular metabolites for untreated MG1655 and Δ*atpA* cells were extracted and quantified on a Q Exactive mass spectrometer with coupled UPLC (Thermo Scientific; Waltham, MA) as described above without the use of spike-in standards. Extracted metabolites were filtered using Phree phospholipid removal filters (Phenomenex; Torrence, CA). All raw metabolomics data are provided in Supplemental Tables 1 and 5.

### Bacterial growth kinetics

Overnight cultures of cells grown in MOPS rich were inoculated 1:20,000 into fresh media. 200 μL of diluted cultures were dispensed into 96-well clear round bottom microplates. Microplates were incubated at 37°C in a Tecan Infinite M Plex microplate reader at 900 rpm. OD_600_ was measured every 12.5 minutes for 12 hours.

### Oxygen consumption and extracellular acidification quantification

Oxygen consumption and extracellular acidification rates were measured using an Agilent Seahorse XF^e^96 Extracellular Flux Analyzer (Agilent; Santa Clara, CA) as described previously^18,22,23^. Briefly, Seahorse XF Pro cell culture microplates were pre-coated with 15 μL 100 μg/mL poly-D-lysine. Overnight cultures grown in MOPS rich media were inoculated 1:500 into fresh media. Cells were grown until they reached OD_600_ ≥ 0.1 and then back-diluted to OD_600_ = 0.0025. 200 μL of diluted cells were dispensed into coated microplates, and microplates were centrifuged for 10 minutes at 1,500 rcf to adhere cells. Oxygen consumption rate (OCR) and extracellular acidification rate (ECAR) measurements were made at 4-minute intervals (2.5 minutes for measurement and 1.5 minutes for mixing) for 4 cycles. Reported values are derived from 5 technical replicates for each biological replicate. Samples were randomized on each plate to control for potential systematic biases. For experiments involving piceatannol, cells were treated with 200 μM piceatannol or 1% DMSO after cultures were back-diluted to OD_600_ = 0.0025. Measurements were taken as described above.

### Membrane potential quantification

DiOC_2_(3) (Thermo Scientific) exhibits green fluorescence in all bacterial cells at low concentrations, but dye import increases with increasing membrane potential, causing the dye to self-associate and for peak fluorescence emissions to shift from 530 nm to 670 nm. Overnight cultures grown in MOPS rich media were inoculated 1:500 into fresh media and grown until OD_600_ ≥ 0.1. Cultures were back-diluted to OD_600_ = 0.1 and treated with 30 μM DiOC_2_(3) and 1 mM EDTA. Treated cells were incubated in the dark at 37°C for 1 hour. 1 μL of treated culture was placed atop a 1.5% agarose pad (Thermo Scientific), mounted on a slide, and sealed with a coverslip. Cells were imaged on a Nikon Ti2 Eclipse microscope (Nikon; Melville, NY), using a 100x oil objective. Cells were excited at 488 nm and emissions were measured at 510-531 nm and 590-624 nm. Fluorescence emissions for each cell were quantified using the MicrobeJ plugin^86^ of ImageJ to segment cells. Fluorescence measurements were background subtracted and DiOC_2_(3) fluorescence ratios were calculated for each cell. Reported values are derived from the median of ≥ 135 cells per biological replicate.

### Minimum inhibitory concentration (MIC) determination

MICs for ciprofloxacin and piceatannol were measured by microbroth dilution. Ciprofloxacin or piceatannol was 1.5-fold serially diluted in MOPS rich media at 100 μL volumes in a 96-well round bottom microplate. The highest working concentration was 40 ng/mL for ciprofloxacin and 100 μg/mL for piceatannol. The last 2 columns of each microplate contained drug-free controls. Overnight cultures grown in MOPS rich media were inoculated 1:10,000 into fresh media and 100 μL of diluted cultures were added to each well of the ciprofloxacin- or piceatannol-loaded microplates. The last column was not inoculated with cells as a cell-free control. Plates were sealed with Breathe-Easy permeable membranes (Sigma-Aldrich) and incubated for 22 hours. OD_600_ was measured on an Tecan Infinite M Plex microplate reader.

### Resistance evolution

Resistance evolution experiments were performed by serial-dilution passaging. 1 mg/mL stocks of ciprofloxacin were prepared and stored at -20°C for consistent chemical preparation. Aliquots were thawed and diluted for use in each evolution cycle and were discarded afterwards. Ciprofloxacin-loaded microplates were prepared above as for MIC experiments. Evolution experiments were initiated by growing ancestral *E. coli* cells in MOPS rich media and inoculating 1:2,500 into fresh media. For each cycle diluted cultures were dispensed into each ciprofloxacin-loaded plate to achieve a final inoculum of 1:5,000 (∼1-2·10^6^ colony forming units) Plates were sealed with Breathe-Easy permeable membranes and incubated for 22 hours. OD_600_ was then measured on an Tecan Infinite M Plex microplate reader. The MIC_50_ was calculated as the concentration of ciprofloxacin required to inhibit growth by ≥50%, compared to ciprofloxacin-free growth controls. At each subsequent evolution cycle, the culture from the highest ciprofloxacin concentration in which bacteria grew (OD_600_ ≥ 0.1) was diluted 1:2,500 and re-inoculated into freshly prepared ciprofloxacin-loaded microplates at a volume of 100 μL to achieve 1:5,000 final dilution. Bacteria were passaged for a total of 8 cycles. Fold change MIC_50_ was calculated by dividing by the MIC_50_ of each cycle by the MIC_50_ of Cycle 1. For experiments involving biochemical supplementation, 12.5 μM piceatannol or 15 U/mL catalase was included in the growth media throughout the entire experiment.

### Time-kill kinetics

Ciprofloxacin time-kill experiments were performed as previously described^18^. Briefly, overnight cultures grown in MOPS rich were inoculated 1:500 into fresh media and grown to OD_600_ ≥ 0.1. Cultures were back-diluted to OD_600_ = 0.1 and treated with 16 ng/mL or 18 ng/mL ciprofloxacin as appropriate for achieving ∼3-log reduction in survival 4 hours post-treatment (Extended Data Fig. 7). Hourly samples were collected and serially diluted in PBS for colony enumeration 24 hours later. Piceatannol or catalase supplementation experiments were performed in the presence of 100 μM piceatannol, 15 U/mL catalase, or 1% DMSO. Bioenergetic stress manipulation experiments involving media switching were performed by preconditioning cells in MOPS rich or MOPS minimal media, pelletting by centrifugation at 6,000 rcf for 2 minutes, washing twice, and then resuspending in the target working media (MOPS rich or MOPS minimal) at OD_600_ = 0.1 in the presence of ciprofloxacin.

### RNA sequncing and analysis

Overnight cultures grown in MOPS rich media were inoculated 1:500 into fresh media and grown to OD_600_ ≥ 0.1. 600 μL cells were then mixed with 1200 μL RNAprotect Bacteria Reagent (Qiagen; Germantown, MD) to inactivate RNase activity. Total RNA was extracted using RNeasy Micro Kits (Qiagen). RNA concentrations and RNA integrity were measured on an Agilent 4200 Tapestation. Ribosomal RNA was depleted using the Ribo-Zero Plus rRNA Depletion Kit (Illumina; San Diego, CA). cDNA libraries were prepared using the NEBNext Ultra II Directional RNA Library Prep Kit for Illumina (New England Biolabs; Ipswich, MA). RNA sequencing was performed on an Illumina NovaSeq6000 system with 100x coverage.

Raw sequencing reads were aligned to the U00096.3 NCBI *E. coli* MG1655 reference genome by Bowtie 2^87^. Read counts were compiled using featureCounts^88^ and quantile normalized by qsmooth^89^. Data quality, adapter and quality trimming statistics, and alignment and count metrics were compiled and assessed using MultiQC^90^. Differential gene expression analysis was performed using DESeq2^91^. Differentially expressed genes were defined to possess false discovery rate (FDR)-corrected p-values ≤ 0.05 and log_2_ fold changes ≥ 1 or ≤ -1. Gene Ontology (GO) analyses of differentially expressed genes were performed using PANTHER^92^. FDR correction was performed using the Benjamini-Hochberg method^93^. Normalized sequencing counts, differential expression analyses, and GO enrichments are available in Supplemental Tables 2 and 3.

### Genome-scale metabolic modeling

Genome-scale metabolic modeling was performed using the iML1515 model of *E. coli* metabolism^48^ as described previously^18^. pEmpty cells were modeled using the default iML1515 model. pF_1_ cells were modeled by adding an ATP sink reaction (ATPASE) defined as “atp_c + h2o_c → adp_c + h_c + pi_c”. The lower bound for the ATP sink reaction was assigned at 1.25 to guarantee ATP hydrolysis in the pF_1_ model. pNOX cells were modeled by adding a NADH oxidase reaction (NOX) defined as “h_c + nadh_c + o2_c → h2o2_c + nad_c”. The lower bound for the NADH oxidase reaction was assigned at 1.33 to guarantee NADH hydrolysis in the pNOX model. Normalized RNA sequencing counts were applied as modeling constraints to iML1515 using the iMAT algorithm^46,47^. Flux variability analysis^49^ was performed using the COBRApy toolbox^94^ with 10,000 flux samples collected for each model by optGpSampler^95^. Model simulations are summarized in Supplemental Table 4.

### H_2_O_2_ production

H_2_O_2_ production was quantified using a highly specific horseradish peroxidase and Amplex UltraRed assay^50^. Overnight cultures grown in MOPS rich media containing 10 U/mL catalase were inoculated 1:500 into fresh media containing catalase and grown to OD_600_ ≥ 0.1. Cells were then pelleted by centrifugation at 6,000 rcf for 2.5 minutes, washed 3 times with fresh media without catalase, and resuspended in fresh media without catalase. Washed cultures were back-diluted to OD_600_ = 0.05. 100 μL diluted culture was dispensed into wells of a 96-well black, clear-bottom microplate. A standard curve for H_2_O_2_ was generated by performing 2-fold serial dilutions at 100 μL volumes. 50 μL of MOPS rich media containing 200 μM Amplex UltraRed was dispensed into each well of the microplate, followed immediately by 50 μL of MOPS rich media containing 100 μg/mL horseradish peroxidase. OD_600_ and fluorescence emissions (550-590 nm) were measured on a Tecan Infinite M-plex microplate reader at 15 minute intervals with 488 nm excitation and 900 rpm shaking and 37°C incubation between reads. Absolute H_2_O_2_ concentrations for each measurement were computed from the standard curve and H_2_O_2_ concentrations were normalized to cell density as measured by OD_600_.

### Carbonylated protein quantification

Protein carbonylation was quantified using the OxiSelect Protein Carbonyl ELISA kit (Cell Biolabs; San Diego, CA) according to the manufacturer’s instructions. Briefly, overnight cultures grown in MOPS rich media were inoculated 1:500 into fresh media and grown to OD_600_ ≥ 0.1. Aliquots with biomass equivalents to 25 mL of cell culture at OD_600_ = 0.1 were pelleted by centrifugation at 4,250 rcf for 10 minutes, washed 3 times with ice-cold PBS, flash frozen, and stored at -80°C before extraction. Cell pellets were lysed by addition with 200 μL Bacterial Protein Extraction Reagent (Thermo Scientific), 100 μg/mL lysozyme, and 5 U/mL DNase I. Total protein abundance of each sample lysate was quantified using a Pierce BCA Protein Assay Kit (Thermo Scientific) according to the manufacturer’s instructions. Samples were diluted to equal protein concentrations prior to performing the OxiSelect ELISA assay. Absorbance measurements were taken at 450 nm on an Agilent Biotek Synergy H1 microplate reader.

### Oxidized deoxyguanosine quantification

Oxidized deoxyguanosine (8-oxo-dG)) was quantified using the OxiSelect Oxidative DNA Damage ELISA kit (Cell Biolabs) according to the manufacturer’s instructions. Frozen bacterial pellets were obtained as described above for carbonylated protein quantification experiments. Total DNA was extracted using the DNeasy UltraClean Microbial Kit (Qiagen) according to the manufacturer’s instructions. DNA concentrations were measured using a NanoDrop One Microvolume UV-Vis Spectrophotometer (Thermo Scientific). Samples were diluted to equal DNA concentrations. Double strand DNA was converted to single strand DNA by incubating samples at 95°C for 5 minutes and rapidly chilling on ice. Single strand DNA was digested with 10 units of Nuclease P1 before performing 8-oxo-dG measurements using the OxiSelect ELISA kit. Absorbance measurements were taken at 450 nm on an Agilent Biotek Synergy H1 microplate reader.

### Mutation rate estimation

Mutation rates were measured using Luria-Delbruck fluctuation assays^53^. Overnight cultures grown in MOPS rich media were inoculated 1:250,000 into fresh media. 1 mL of diluted culture (∼10^3^-10^4^ CFUs) was dispensed into the wells of a 96-well, v-bottom, square deep well plate. Plates were sealed using Breathe-Easy permeable membranes and incubated for 24 hours at 37°C and 900 rpm shaking. 500 μL stationary phase cultures were plated on LB agar plates ± 100 μg/mL rifampicin. Colonies were enumerated after 24 hours incubation at 37°C. Rifampicin resistance (Rif^R^) CFUs were normalized by total CFUs. Reported values were scaled by 10^8^.

### (p)ppGpp quantification

Intracellular (p)ppGpp abundance was quantified using the S2 RNA-based fluorescent biosensor^58^. Overnight cultures grown in MOPS rich were inoculated 1:500 into fresh media and grown to OD_600_ ≥ 0.1. Cultures were back-diluted to OD_600_ = 0.1 and 1 mM IPTG was added to induce S2 expression. After 1.5 hours of induction, 50 μL aliquots were added to 50 μL of MOPS rich media containing 1 mM IPTG and 400 μM DFHBI-1T in 1.5 mL microcentrifuge tubes such that the final IPTG and DFHBI-1T concentrations were 1 mM and 400 μM, respectively. DFHBI-1T is the cognate dye for the S2 sensor. Cells were then treated with 18 ng/mL ciprofloxacin or H_2_O control. Tubes were incubated at room temperature in the dark for 1 hour before imaging. 1 μL of cells were placed atop a 1.5% agarose pad mounted on a slide and sealed with a coverslip. Cells were imaged on a Nikon Ti2 Eclipse microscope, using a 100x oil objective. Fluorescence measurements were taken at 488 nm and 510-531 nm emission. Background-subtracted single-cell fluorescence intensities were quantified using the MicrobeJ^86^ plugin of ImageJ. Reported values are derived from the median fluorescence of n ≥ 100 cells per replicate.

### Statistical analyses

Statistical significance testing was performed in Prism v10 (Graphpad; San Diego, CA). Normality testing was performed using the Shapiro-Wilk test. Homogeneity of variance testing was performed using the F test or the Bartlett test, where appropriate. All two-sample comparisons were performed using the Mann-Whitney test or unpaired *t*-test. All multiple-sample comparisons were performed by one-way ANOVA with the Dunnett’s or Šídák’s multiple comparison test; Welch’s ANOVA with the Dunn’s multiple comparison test; or two-way ANOVA with the Šídák’s multiple comparison test. Selection of parametric vs. non-parametric tests was determined by tests for normality and homogeneity of variance.

## ACKNOWLEDGEMENTS

We thank the Rutgers Metabolomics Core and Genomics Center for support in collecting metabolomics and performing RNA sequencing measurements, respectively. We thank Douglas McCloskey (BioMed X) for assistance with metabolomics experiments. We thank James Imlay (UIUC) for helpful discussions.

## AUTHOR CONTRIBUTIONS

B.L. and J.H.Y conceptualized the study. B.L., S.S., M.S., G.M., M.G, A.S., N.O., T.C., N.C., and J.H.Y. executed experiments. B.L. and J.H.Y. analyzed and visualized the data and wrote the manuscript. J.H.Y. supervised the study.

## DATA AND MATERIALS AVAILABILITY

All data are available in the main text or supplementary materials. RNA sequencing data is available in on the Gene Expression Omnibus (GEOXXXXXX) and Sequence Read Archive (SRAXXXXXX). Analysis code is deposited on GitHub (https://github.com/jasonhyang).

---

**Extended Data Fig. 1.**
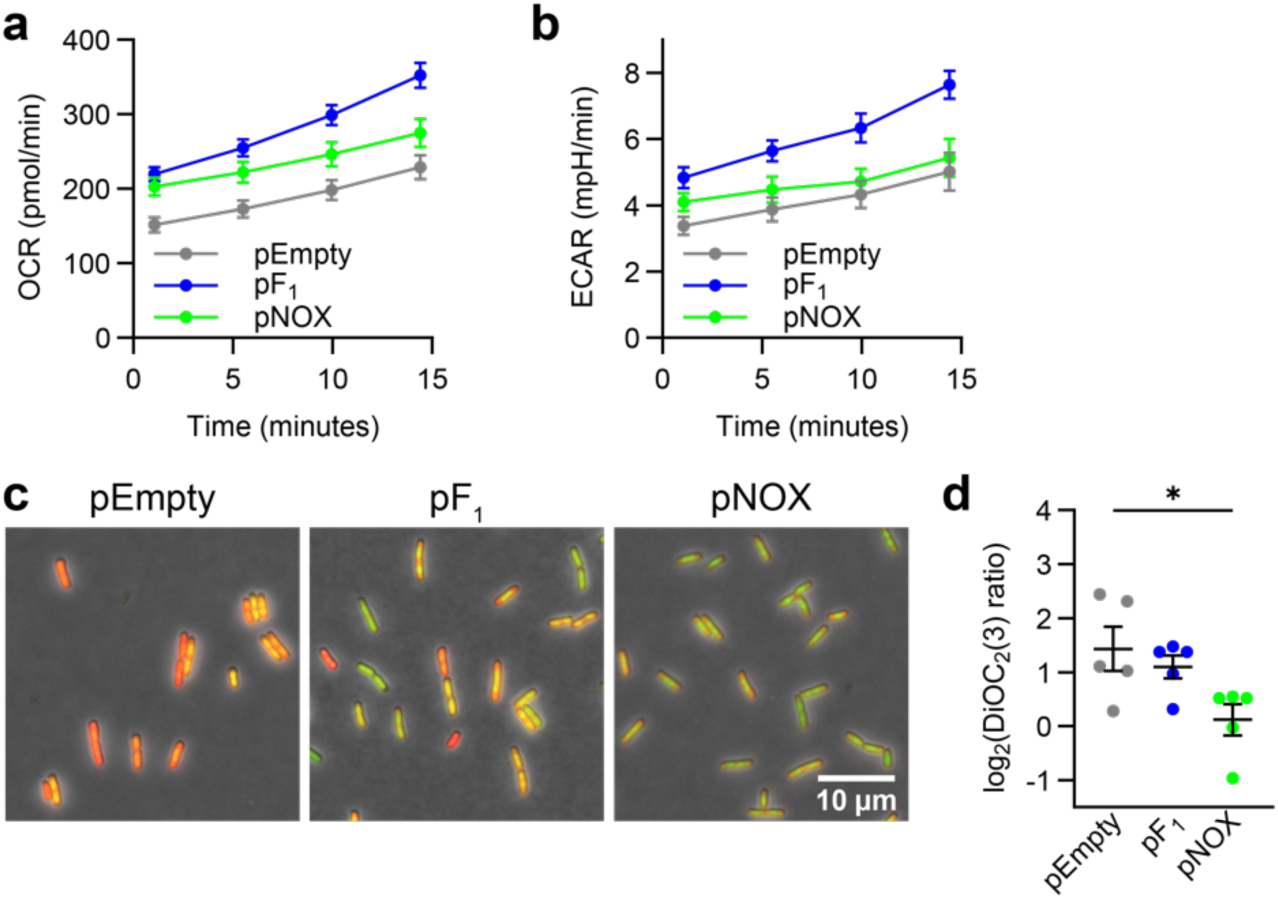
Bioenergetic stress alters respiratory and glycolytic activity. **a**, Oxygen consumption rates (OCR) as a reporter of respiratory activity for exponential phase pEmpty, pF_1_, and pNOX cells (n = 4). Data depicted as mean ± SEM. **b**, Extracellular acidification rates (ECAR) as a reporter of glycolytic activity (n = 4). Data depicted as mean ± SEM. **c**, Representative fluorescence microscopy images of exponential phase pEmpty, pF_1_, and pNOX cells labelled with the membrane potential-sensitive dye DiOC_2_(3) for 1 hour. **d**, Single-cell DiOC_2_(3) fluorescence ratios for exponential phase pEmpty, pF_1_, and pNOX cells. Decreased ratio indicates membrane depolarization. Data points depict median DiOC_2_(3) ratio per replicate (n ≥ 135 cells per replicate from 5 biological replicates). Error bars depict grand means ± SEM. All experiments performed in MOPS rich media. ns > 0.05, \**p* ≤ 0.05, \*\*\*\**p* ≤ 0.0001. Non-significant comparisons not shown.

**Extended Data Fig. 2.**
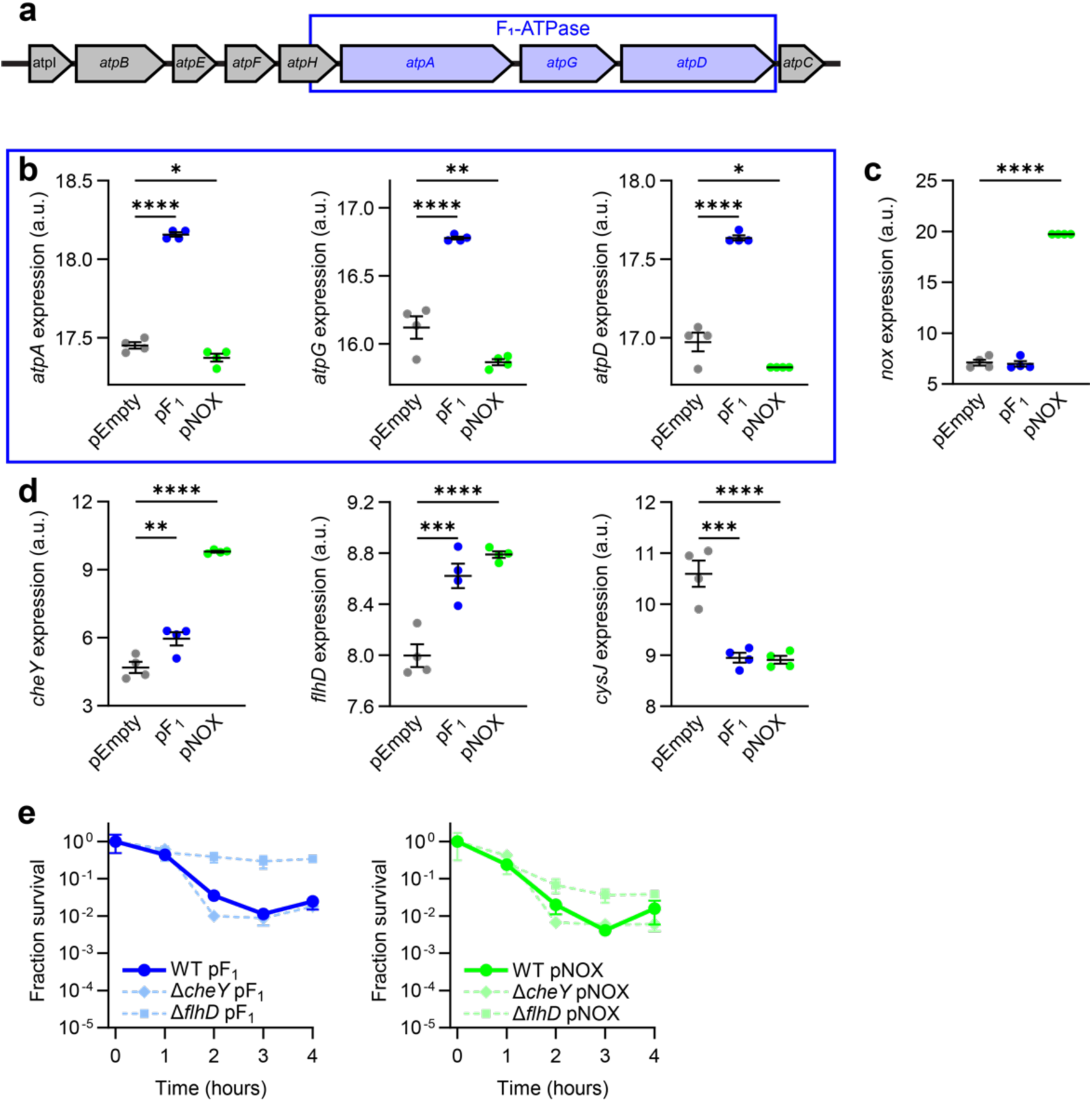
Bioenergetic stress-induced persistence is not caused by increased motility. **a**, Schematic of the *E. coli* ATP synthase operon. F_1_-ATPase genes (*atpAGD*) are outlined in blue. **b**, *atpAGD* expression for exponential phase pEmpty, pF_1_, and pNOX cells. **c**, *Streptococcus pneumoniae nox* expression for exponential phase pEmpty, pF_1_, and pNOX cells. **d**, *cheY* and *flhD* expression for exponential phase pEmpty, pF_1_, and pNOX cells. Expression data depicted as quantile normalized log_2_-transformed transcripts per million counts as measured by RNA-sequencing. Error bars depict mean ± SEM. **e**, Ciprofloxacin lethality following treatment with 18 ng/mL ciprofloxacin for Δ*cheY* or Δ*flhD* cells expressing pEmpty, pF_1_, or pNOX. Data reported as change in colony forming units (CFUs) relative to time 0 for all time-kill experiments. Data depicted as mean ± 95% CI. All experiments performed in MOPS rich media. \**p* ≤ 0.05, \*\**p* ≤ 0.01, \*\*\*\**p* ≤ 0.0001. Non-significant comparisons not shown.

**Extended Data Fig. 3.**
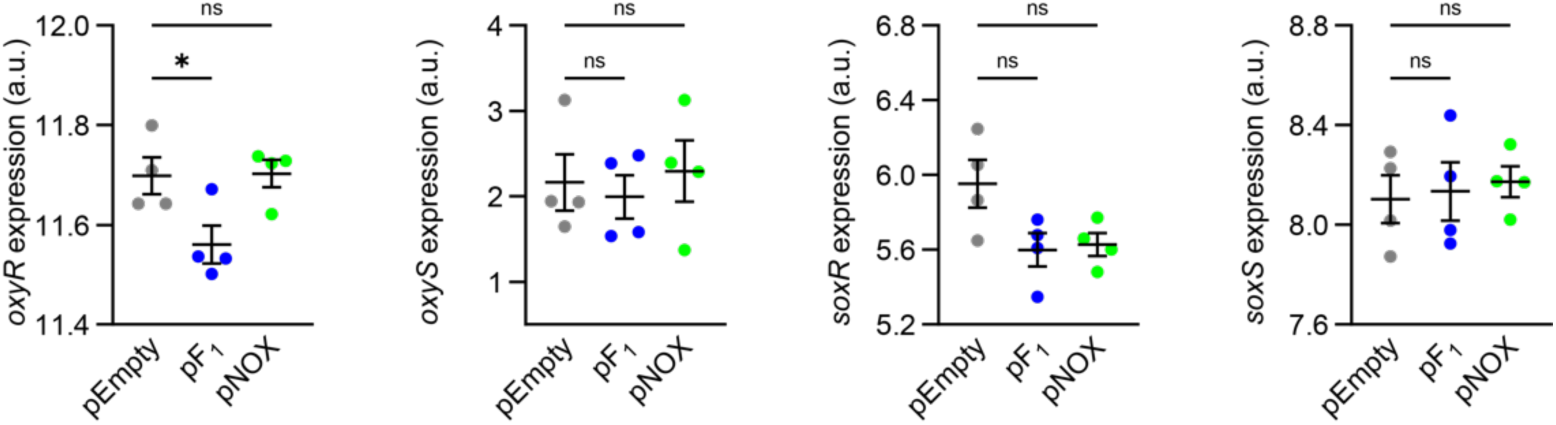
Bioenergetic stress does not significantly induce the OxyR or SoxRS stress responses. *oxyR*, *oxyS*, *soxR*, and *soxS* expression for exponential phase pEmpty, pF_1_, and pNOX cells. Data depicted as quantile normalized log_2_-transformed transcripts per million counts as measured by RNA-sequencing. Error bars depict mean ± SEM. ns > 0.05, \**p* ≤ 0.05.

**Extended Data Fig. 4.**
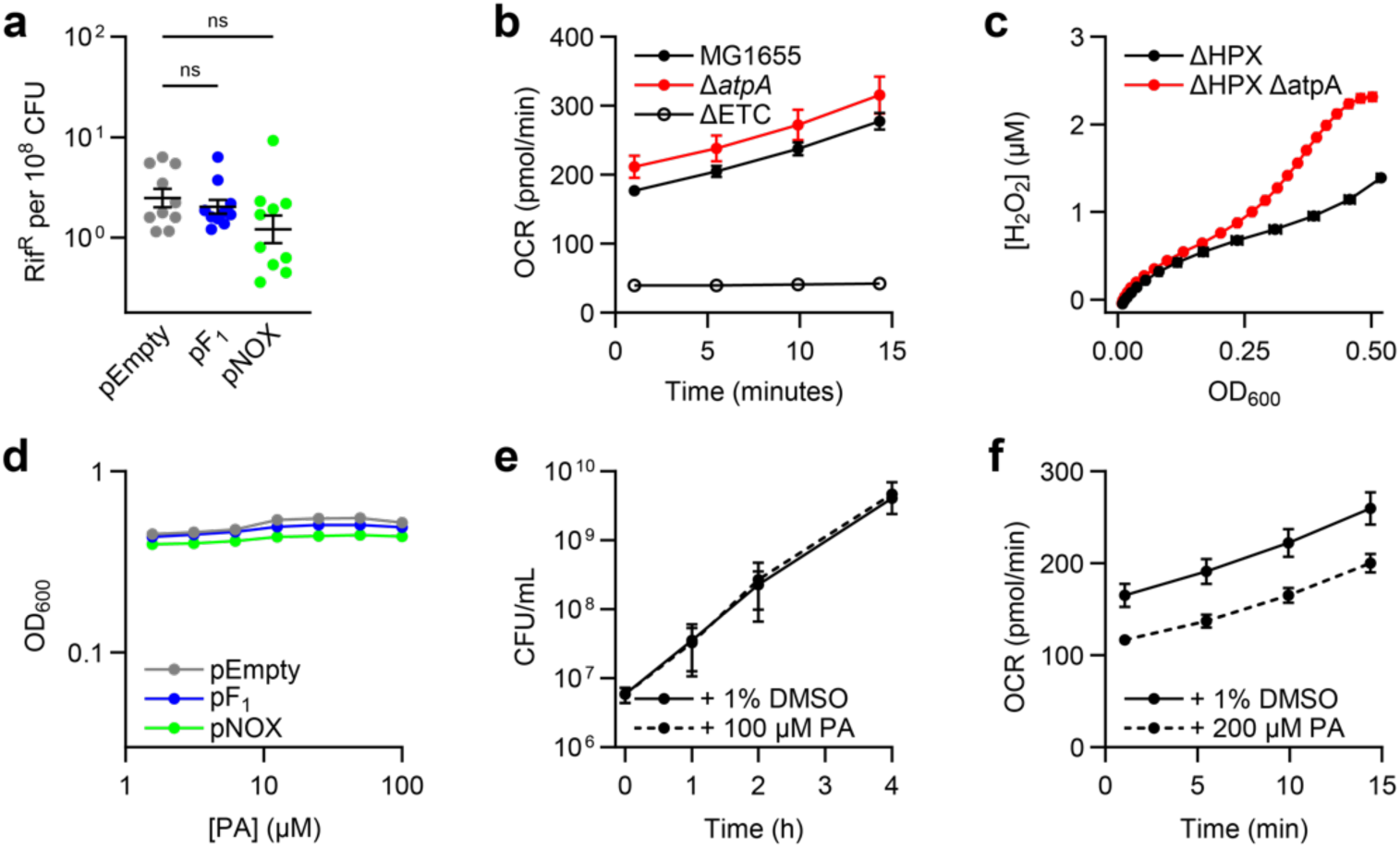
Bioenergetic stress accelerates antimicrobial resistance evolution via stress-induced mutagenesis and increased respiration. **a**, Basal mutation rates do not differ between pEmpty, pF_1_, and pNOX cells as determined by Luria-Delbrück fluctuation assays^53,54^ (n ≥ 10). **b**, Oxygen consumption rates (OCR) as a reporter of respiratory activity for exponential phase wild-type MG1655, Δ*atpA*, and *Δ*ETC cells. **c**, Integrated H_2_O_2_ production by ΔHPX and ΔHPX Δ*atpA* cells. **d**, Growth measurements in the presence of piceattanol (PA) after 1:20,000 dilution from stationary phase. Piceattanol does not inhibit growth of pEmpty, pF_1_, or pNOX cells. **e**, Growth of MG1655 cells ± treatment with 100 µM piceattanol, as determined by CFUs. Piceattanol does not impose growth defects in MG1655 cells. **f**, OCR for exponential phase MG1655 cells with and without treatment with 100 µM piceattanol. Piceattanol inhibits cellular respiration. All experiments performed in MOPS rich media. n = 4 for all experiments unless otherwise indicated. Error bars depict mean ± SEM unless otherwise indicated. ns > 0.05.

**Extended Data Fig. 5.**
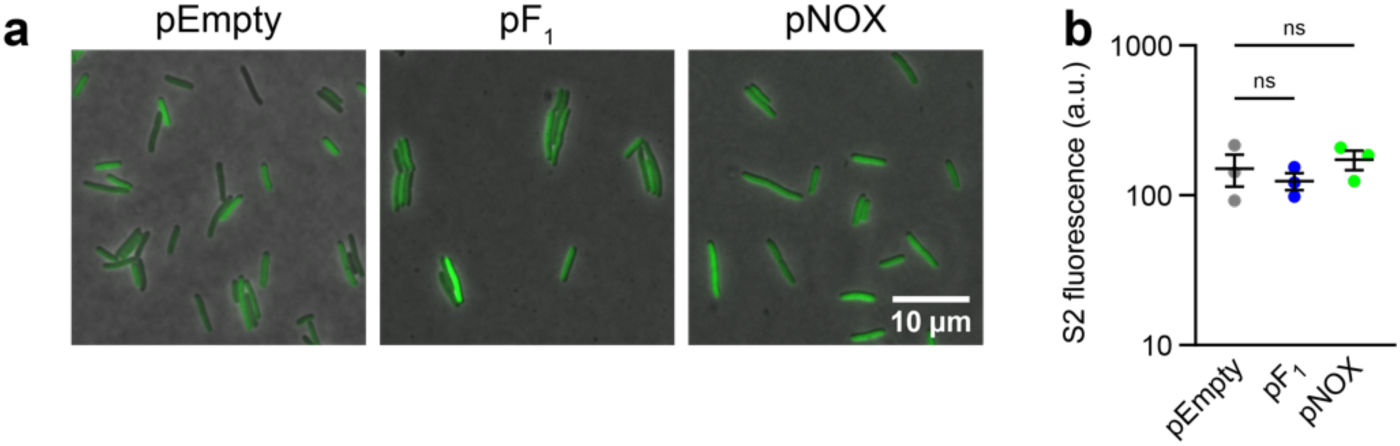
Bioenergetic stress does not basally induce the stringent response. **a**, Representative fluorescence microscopy images of exponential phase *E. coli* BL21(DE3) cells expressing the S2 (p)ppGpp fluorescent biosensor^58^ and pEmpty, pF_1_, or pNOX. **b**, S2 fluorescence quantification (n = 3). Data points depict medians of mean cell intensities. Error bars depict grand mean ± SEM from n ≥ 100 cells per replicate. S2 experiments performed in MOPS rich media. ns > 0.05.

**Extended Data Fig. 6.**
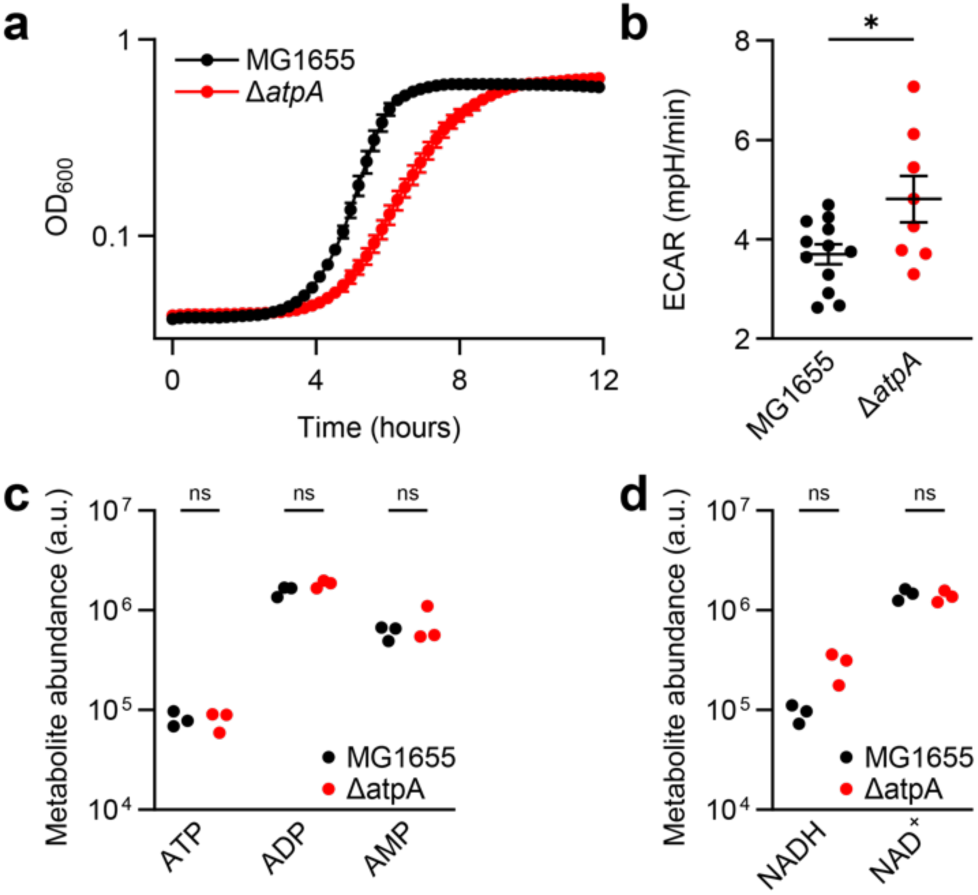
Δ*atpA* cells are not bioenergetically stressed. **a**, Growth curves for MG1655 and Δ*atpA* cells grown in MOPS rich media. **b**, Extracellular acidification rates (ECAR) as a reporter of glycolytic activity for exponential phase MG1655 or Δ*atpA* cells (n = 4; unpaired *t*-test). **c**, ATP, ADP, and AMP abundances for exponential phase wild-type MG1655 and Δ*atpA* cells grown in MOPS minimal media as determined by LC-MS/MS. **d**, NADH and NAD^+^ abundances for exponential phase MG1655 and Δ*atpA* cells. ns > 0.05, \**p* ≤ 0.05.

**Extended Data Fig. 7.**
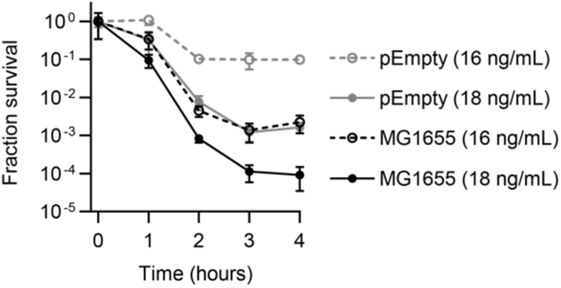
pEmpty cells are less susceptible to ciprofloxacin than MG1655 cells. Ciprofloxacin lethality for MG1655 or pEmpty cells treated with 16 or 18 ng/mL ciprofloxacin in MOPS rich media (n = 4). 18 ng/mL ciprofloxacin elicits similar lethality in pEmpty cells to 16 ng/mL ciprofloxacin in MG1655 cells. Data reported as change in colony forming units (CFUs) relative to time 0. Data depicted as mean ± 95% CI.

**Supplemental Table 1.** Metabolomic profiles for ciprofloxacin-treated cells.

**Supplemental Table 2.** Normalized RNA sequencing counts and differential gene expression analyses.

**Supplemental Table 3.** Gene Ontology analyses.

**Supplemental Table 4.** Genome-scale metabolic modeling analyses.

**Supplemental Table 5.** Metabolomic profiles for Δ*atpA* cells.

**Supplemental Table 6.** Strains and plasmids used in this study.

